# Selfee: Self-supervised Features Extraction of animal behaviors

**DOI:** 10.1101/2021.12.24.474120

**Authors:** Yinjun Jia, Shuaishuai Li, Xuan Guo, Junqiang Hu, Xiao-Hong Xu, Wei Zhang

## Abstract

Fast and accurately characterizing animal behaviors is crucial for neuroscience research. Deep learning models are efficiently used in laboratories for behavior analysis. However, it has not been achieved to use a fully unsupervised method to extract comprehensive and discriminative features directly from raw behavior video frames for annotation and analysis purposes. Here, we report a self-supervised feature extraction (Selfee) convolutional neural network with multiple downstream applications to process video frames of animal behavior in an end-to-end way. Visualization and classification of the extracted features (Meta-representations) validate that Selfee processes animal behaviors in a comparable way of human understanding. We demonstrate that Meta-representations can be efficiently used to detect anomalous behaviors that are indiscernible to human observation and hint in-depth analysis. Furthermore, time-series analyses of Meta-representations reveal the temporal dynamics of animal behaviors. In conclusion, we present a self-supervised learning approach to extract comprehensive and discriminative features directly from raw video recordings of animal behaviors and demonstrate its potential usage for various downstream applications.

## INTRODUCTION

Extracting representative features of animal behaviors has long been an important strategy to study the relationship between genes, neural circuits, and behaviors. Traditionally, human observations and descriptions are the primary solutions for animal behavior analysis^1^. Well-trained researchers would define a set of behavior patterns and compare their intensity or proportion between experimental and control groups. With the emergence and thrive of machine learning methodology, supervised learning has been assisting human annotations and achieved impressive results^2-4^. Nevertheless, supervised learning is limited by prior knowledge and manually assigned labels, thus could not identify behavioral features that are not annotated.

Other machine learning methods were then introduced to the field which were designed to extract representative features beyond human-defined labels. These methods can be generally divided into two major categories: one estimates animal postures with a group of pre-defined key points of the body parts, and the other directly transforms raw images. The former category marks representative key points of animal bodies, including limbs, joints, trunks, and/or other body parts of interest^5-7^. Those features are usually sufficient to represent animal behaviors. However, it has been demonstrated that the key points generated by pose estimation are less efficient for direct behavior classification or two- dimensional visualization^8,9^. Sophisticated post-processing like recurrent neural networks (RNNs)^8^, non-locomotor movement decomposition^10^, or feature engineerings^9^ can be applied to transform the key points into higher-level discriminative features. Additionally, neglected body parts could be catastrophic. For example, the position of the proboscis of a fly is commonly neglected in behavior studies^9,11^. Still, it is crucial for feeding^12^, licking behavior during courtship^13^, and hardness detection for a substrate^14^. Finally, best to our knowledge, there is no demonstration of these pose-estimation methods applied to multiple animals of the same color with intensive interactions. Thus, the application of pose-estimation to mating behaviors of two black mice, a broadly adopted behavior pradigm^15-17^, could be limited because labeling body parts during mice mounting is challenging even for humans (see Discussion for more details). Therefore, using these feature extraction methods requires rigorously controlled experimental settings, additional feature engineering, and considerable prior knowledge of particular behaviors.

In contrast, the other category transforms pixel-level information, thus retaining more details and requiring less prior knowledge. Feature extraction of images could be achieved by wavelet transforms^18^ or Radon transforms^19^ followed by principal component analysis (PCA), and these transforms can be applied to either 2D images or depth images. However, preprocessing such as segmentation and/or registration of the images is usually required to achieve spatial invariance, a task that is particularly difficult for multi-agent videos. Additionally, these methods usually use fixed transforms and could not be adapted to different behaviors. Flourished deep learning methods, especially convolutional neural networks^20^ (CNNs), could be adaptive to extract features from diversified datasets. Also, they have been proven more potent than classic computer vision algorithms like wavelet transforms^21^ and Radon transforms^22^ on a famous grayscale dataset MNIST, even without supervising^23^. Therefore, we attempt to adopt CNNs to achieve end-to-end feature extractions animal behaviors that are comprehensive and discriminative.

The cutting-edge self-supervised deep learning methods aim to extract representative features for downstream missions by comparing different augmentations of the same image and/or different images^24-28^. Compared with previous techniques, these methods have three major advantages. Firstly, self-supervised or unsupervised methods could completely avoid human biases. Secondly, the augmentations used to create positive samples promise invariance of the neural networks to object sizes, spatial orientations, and ambient laminations so that registration or other preprocessing is not required. Finally, the networks are optimized to export similar results for positive samples and separate negative ones, such that the extracted features are inherently discriminative. Even without negative samples, the networks can utilize differential information within batches to obtain remarkable results on downstream missions like classification or image segmentation^27,29,30^. These advances in self-supervised learning provide a promising way to analyze animal behaviors.

In this work, we develop Selfee (**<u>S el</u>**f -supervised **<u>Fe</u>**atures **<u>E</u>** x traction) that adopts cutting-edge self-supervised learning algorithms and CNNs to analyze animal behaviors. Selfee is trained on massive unlabeled behavior video frames to avoid human bias on annotating animal behaviors, and it could capture a global character of animal behaviors even when detailed postures are hard to see, just like human observation. During the training process, Selfee learns to project images to a low-dimensional space without being affected by shooting conditions, image translation, and rotation, where cosine distance is proper to measure the similarities of original pictures. Selfee also provides potentials for various downstream analyses. We demonstrate that the extracted features are suitable for t-SNE visualization, k-NN-based classification, k-NN-based anomaly detection, and dynamic time warping (DTW). We also show that further integrated modeling, like the autoregressive hidden Markov model (AR-HMM), is compatible with Selfee extracted Meta-representations. After downstream analyses, Selfee provides comparable results with manual annotations on fly behavior like courtship index. We apply Selfee to fruit flies, mice, and rats, three widely used model animals, and validate our results with manual annotations. Discoveries of behavioral phenotypes in mutant flies by Selfee are proven to have biological significance. The performance of Selfee on these model species indicates its potential usage for behavioral studies of non-model animals as well as other tasks. We also provide an open-source Python package and pre-trained models of flies and mice to the community (see more in Code Availability).

## RESULTS

### Workflow of Selfee and its downstream analyses

Selfee is trained to generate Meta-representations at the frame level, which are then analyzed at different time scales. First, grayscale videos are decomposed into single frames, and three tandem frames are stacked into a live-frame to generate a motion- colored RGB picture (Figure 1A). These live-frames preserve not only spatial information (e.g., postures of each individual or relative distances and angles between individuals) within each channel but also temporal information across different channels. Live-frames are used to train Selfee to produce comprehensive and discriminative representations at the frame level (Figure 1B). These representations can be later used in numerous applications. For example, anomalous detection on mutant animals can discover new phenotypes compared with their genetic controls (Figure 1C). Also, the AR-HMM could be applied to model the micro-dynamics of behaviors, such as the duration of states or the probabilities of state transitions^18^. The AR-HMM splits videos into modules and yields behavioral state usages that visualize differences between genotypes (Figure 1D). In contrast, DTW could compare the long-term dynamics of animal behaviors and capture global differences at the video level^31^ by aligning pairs of time series and calculating their similarities (Figure 1E). These three demonstrations cover different time scales from frame to video level, and other downstream analyses could also be incorporated into the workflow of Selfee.

**Figure 1.**
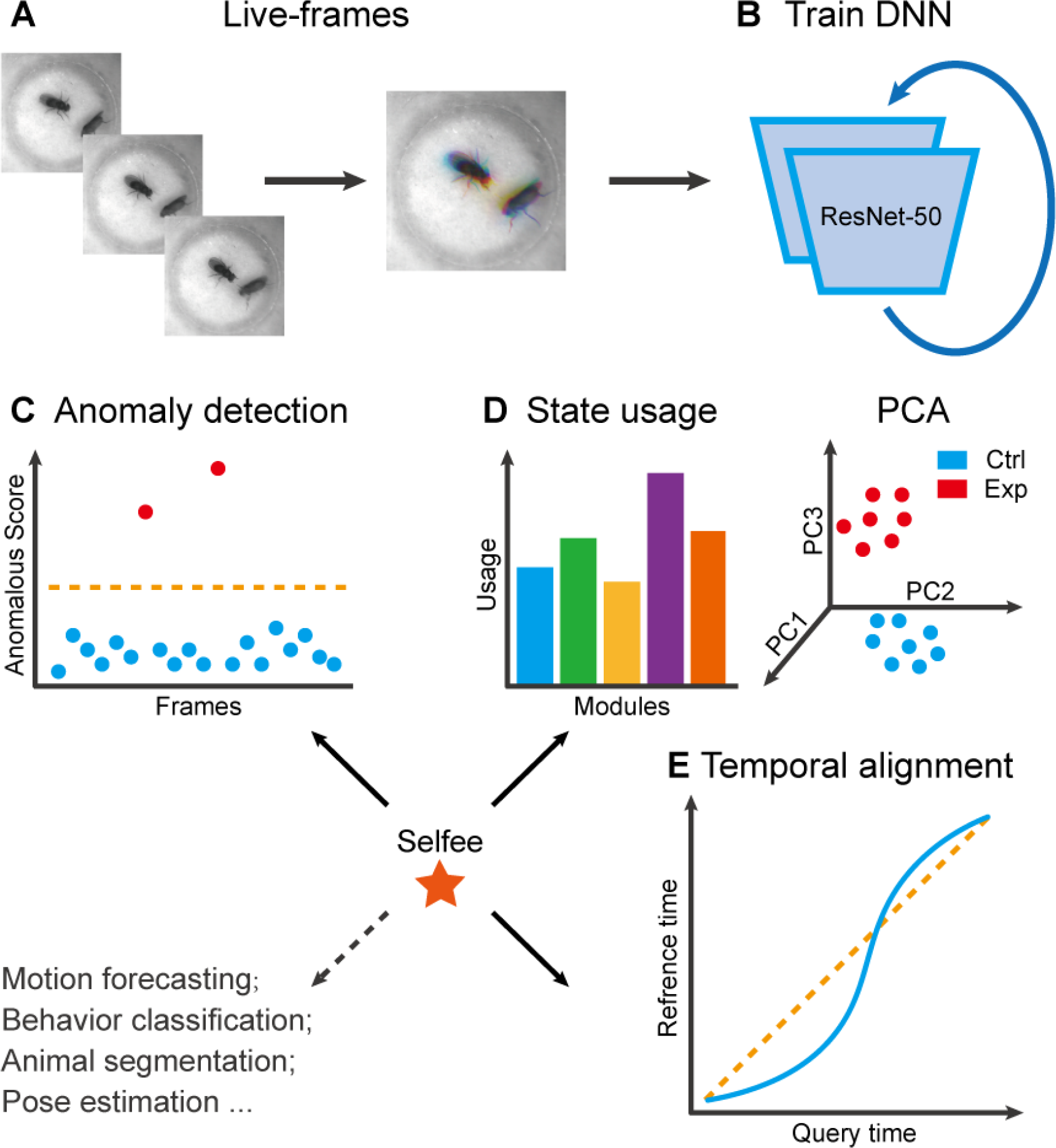
The framework of Selfee and its downstream applications. (A) One live-frame is composed of three tandem frames in R, G, and B channels, respectively. (B) Live-frames are used to train Selfee, which adopts a backbone of ResNet-50. (C, D, and E) Representations produced by Selfee could be used for anomaly detection that could identify unusual animal postures in the query video compared with the reference videos (C) AR-HMM (autoregressive hidden Markov model) that models the local temporal characteristics of behaviors and clusters frames into modules (states) and calculates stages usages of different genotypes (D) DTW (dynamic time warping) that aligns behavior videos to reveal differences of long-term dynamics (E) and other potential tasks including behavior classification, forecasting, or even image segmentation and pose estimation after properly modifying and finetuning of the neural networks.

Compared with previous machine learning frameworks for animal behavior analysis, Selfee has three major advantages. First, Selfee and the Meta-representations could be used for various tasks. The contrastive learning process of Selfee would allow output features to be appropriately compared by cosine similarity. Therefore, distance-based applications, including classification, clustering, and anomaly detection, would be easily realized. It was also reported that with some adjustment of backbones, self-supervised learning would facilitate tasks such as pose estimation^32^ and object segmentation^28,33^. Those findings indicate that Selfee could be generalized, modified, and finetuned for animal pose estimation or segmentation tasks. Second, Selfee is a fully unsupervised method developed to annotate animal behaviors. Although some other techniques also adopt semi-supervised or unsupervised learning, they usually require manually labeled pre-defined key points of the images ^8,10^; some methods also require expert-defined programs for better performance^9^. Key point selection and program incorporation require a significant amount of prior knowledge and are subject to human bias. In contrast, Selfee does not need any prior knowledge. Finally, Selfee is relatively hardware-inexpensive. Training Selfee only takes eight hours on a single RTX 3090, and the inference speed could reach 800 frames per second. Selfee could accept top-view 2D greyscale video frames as inputs so that neither depth cameras^18^ nor fine-calibrated multi-view camera arrays^10^ is required. Therefore, Selfee can be trained and used with routinely collected behavior videos on ordinary desktop workstations, warranting its accessibility by biology laboratories.

### Siamese convolutional neural networks capture discriminative representations of animal posture

Selfee contains a pair of Siamese CNNs trained to generate discriminative representations for live-frames. ResNet-50^34^ is chosen as the backbone whose classifier layer is replaced by a three-layer multi-layer perceptron (MLP). These MLPs are called projectors which yield final representations during the inference stage. There are two branches in Selfee. The main branch is equipped with an additional predictor, while the reference branch is a copy of the main branch (the SimSiam style^29^). Both branches contain group discriminators after projectors and perform dimension reduction on extracted features for online clustering (Figure 2B).

**Figure 2.**
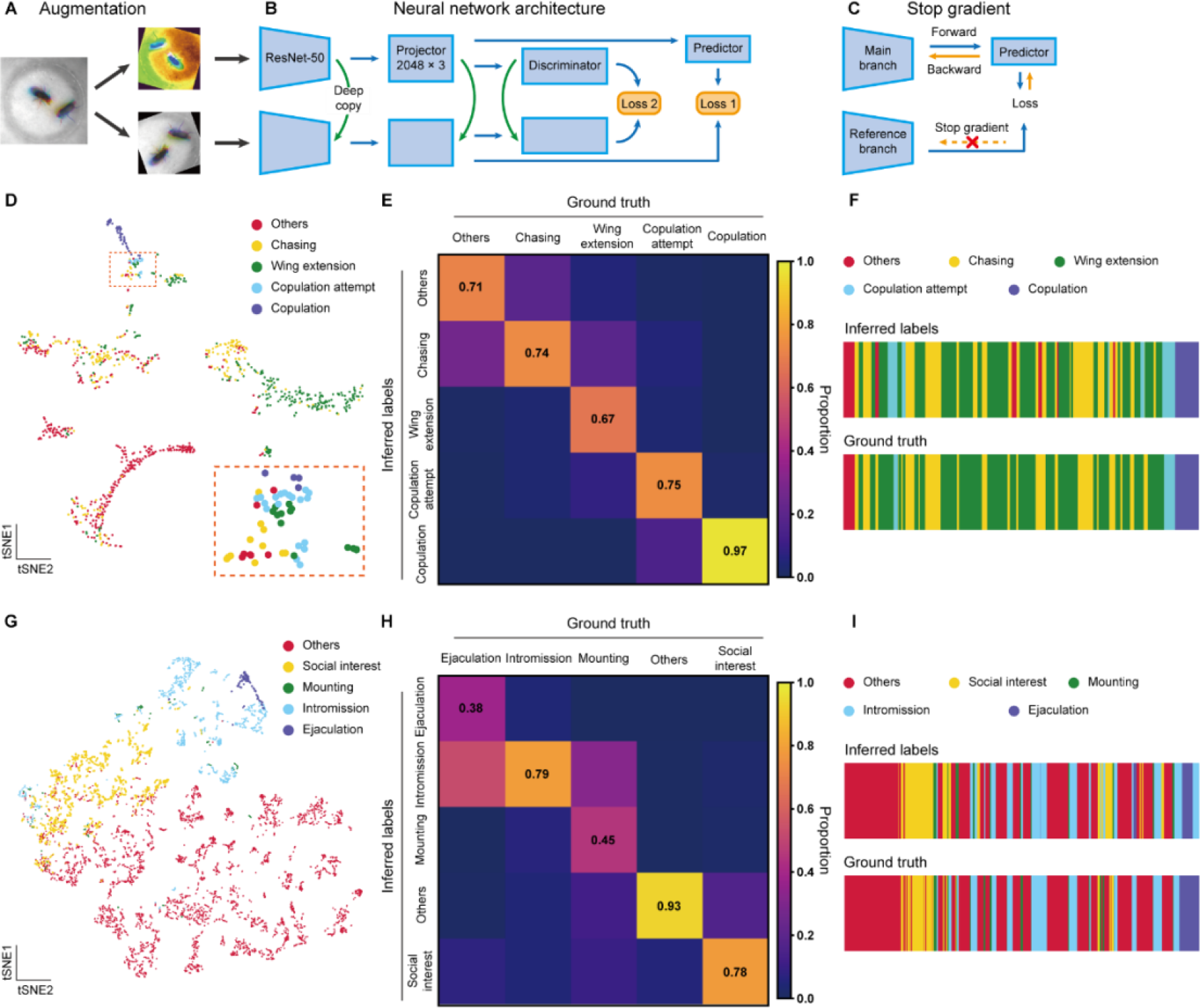
The network structure of Selfee and validation of Selfee with human annotations. (A) The architecture of Selfee networks. Each live-frame is randomly transformed twice before being fed into Selfee. (B) Selfee adopts a SimSiam-style network structure with additional group discriminators. Loss 1 is canonical negative cosine loss, and loss 2 is the newly proposed CLD loss. (C) The reference branch of Selfee is not involved in backward propagation. (D) Visualization of fly courtship live-frames with t-SNE dimension reduction. Each dot was colored based on human annotations. (E) The confusion matrix of the k-NN classifier for fly courtship behavior, normalized by the numbers of each behavior in the ground truth. The average F1 score of the nine-fold cross-validation was 72.4%, and mAP was 75.8%. (F) A visualized comparison of labels produced by the k-NN classifier and human annotations of fly courtship behaviors. (G) Visualization of live-frames of mice mating behaviors with t-SNE dimension reduction. Each dot is colored based on human annotations. (H) The confusion matrix of the LightGBM classifier for mice mating behaviors, normalized by the numbers of each behavior in the ground truth. For the LightGBM classifier, the average F1 score of the nine-fold cross-validation was 67.4%, and mAP was 69.1%. (I) A visualized comparison of labels produced by the LightGBM classifier and human annotations of mice mating behaviors.

During the training stage, batches of live-frames are randomly transformed twice and fed into the main branch and reference branch, respectively. Augmentations applied to live-frames include crop, rotation, flip, and application of the Turbo lookup table^35^ followed by color jitters (Figure 2A, Figure 2—figure supplement 1). The reference branch yields a representation of received frames, while the main branch predicts the outcome of the reference branch. At the same time, they both produce clustering results of the current batch. The main branch is optimized for similar predictions and clustering results as the reference branch, and the reference branch will not receive gradient information to prevent mode collapse^27,29^ (Figure 2C). In this way, Selfee is trained to be invariant to those transforms and focus on critical information to yield discriminative representations.

After the training stage, we evaluated the performance of Selfee with t-SNE visualization and k-NN classification. To investigate whether our model captures human-interpretable features, we manually labeled one clip of *Drosophila* courtship video and visualized those representations with t-SNE dimension reduction. On the t- SNE map, human-annotated courtship behaviors, including chasing, wing extension, copulation attempt, copulation, and non-interactive behaviors (“others”), separated from each other distinctively (Figure 2D).

Meta-representations can also be used for behavior classification. We manually labeled seven 10,000-frame videos (around five minutes each) as a pilot dataset. A weighed k- NN classifier was then constructed as previously reported^24^. Seven-fold cross- validation was performed on the dataset with the k-NN classifier, which achieved a mean F1 score of 72.4% and achieved a similar classification result as human annotations (Figure 2E, F). The classifier had the worst recall score on wing extension behaviors (67% recall), likely because of the ambiguous intermediate states between chasing and wing extension (Figure 2—figure supplement 2A). The precisions also showed that this k-NN classifier tended to have strict criteria with wing extension and copulation and relatively loose criteria with chasing and copulation attempts (Figure 2—figure supplement 2B). It was reported that independent human experts could only reach agreements on around 70% of wing extension frames^36^, comparable to the performance of our k-NN classifier.

We then asked whether Selfee can be generalized to analyze behaviors of other species.

We finetuned fly video pre-trained Selfee with mice mating behavior data. The mating behavior of mice can be defined mainly into five categories^37^, including social interest, mounting, intromission, ejaculation, and others (see Methods for detailed definitions). With t-SNE visualization, we found that five types of behaviors could be separated by Selfee, although mounting behaviors were rare and not concentrated (Figure 2G). We then used eight human-annotated videos to test the k-NN classification performance of Selfee-generated features. We achieved an F1 score of 59.0% (Figure 2—figure supplement 3). Mounting, intromission, and ejaculation share similar static characteristics but are different in temporal dynamics. Therefore, we asked if more temporal information would assist the classification. Using the LightGBM classifier, we achieved a much higher classification performance by incorporating slide moving average and standard division of 81-frame time windows, the main frequencies, and their energy within 81-frame time windows. The average F1 score of eight-fold cross-validation could reach 67.4%, and the classification results of the ensembled classifier (see Methods) were closed to human observations (Figure 2H, I). Nevertheless, it was still difficult to distinguish between mounting, intromission, and ejaculation because mounting and ejaculation are much rarer than social body contact or intromission.

Selfee is more robust than the vanilla SimSiam networks when applied to the behavioral data. Behavioral data often suffer from catastrophic imbalance. For example, copulation attempts are around six-fold rarer than wing extension during fly courtship (Figure 2— figure supplement 5A). Therefore, we added group discriminators to vanilla SimSiam networks which were reported to fight against the long-tail effect proficiently^38^. Aside from overcoming the long-tail effect, we also found group discriminators helpful for preventing mode collapse during ablation studies (Figure 2—figure supplement 5B, C, D, and Supplementary Table 1). Additionally, the convergence can be easily reached on grayscale images of similar objects (two flies), by which CNNs may not be well trained to extract good representations. Applying the Turbo lookup table on grayscale frames brought more complexity and made color jitters more powerful on grayscale images. Selfee would capture more useful features with this Turbo augmentation (Figure 2— figure supplement 5E, F, and Figure 2—figure supplement 6).

### Anomaly detection at the frame level identifies rare behaviors at the sub-second time scale

The representations produced by Selfee could be directedly used for anomaly detection without further post-processing. During the training step, Selfee learns to compare Meta-representations of frames with cosine distance which is also used for anomaly detection. When given two groups of videos, namely the query group and the reference group, the anomaly score of each live-frame in the query group is calculated by two steps (Figure 3A). First, distances between the query live-frame and all reference live- frames are measured, and the k-nearest distance is referred to as its inter-group score (IES). Without further specification, k equals 1 in all anomaly detections in this work.

**Figure 3.**
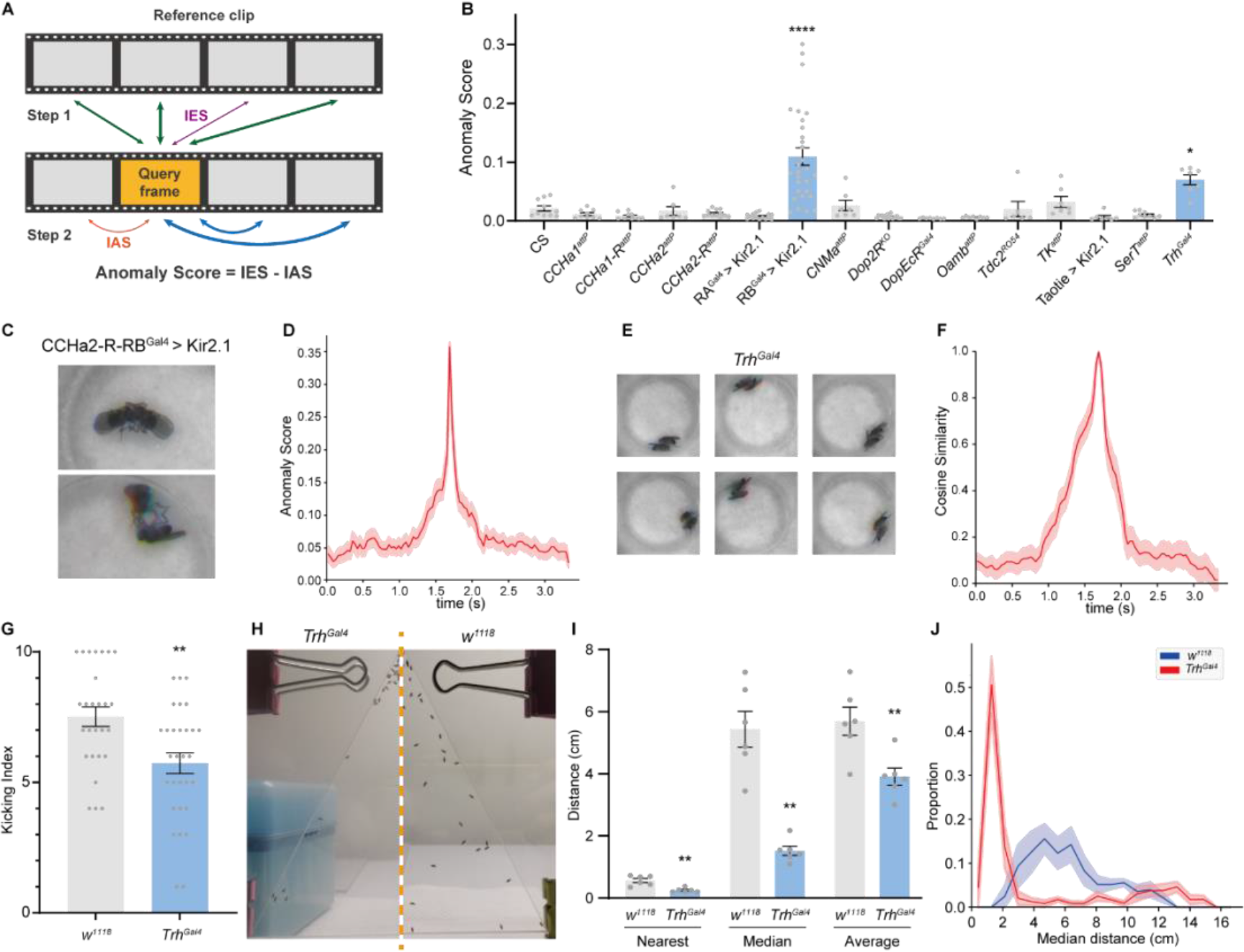
Anomalous posture detection using Selfee-produced features. (A) The calculation process of anomaly scores. Each query frame is compared with every reference frame, and the nearest distance was named IES (the thickness of lines indicates distances). Each query frame is also compared with every query frame, and the nearest distance is called IAS. The final anomaly score of each frame equals IES minus IAS. (B) Genetic screening of fifteen fly lines with mutations in neurotransmitter genes or with specific neurons silenced (n≥6 for each genotype). RA is short for CCHa2-R-RA, and RB is short for CCHa2-R-RB. CCHa2-R-RB^Gal4^ > Kir2.1, q < 0.0001; *Trh^Gal4^*, q = 0.0432; one-way ANOVA with Benjamini and Hochberg correction. (C) Examples of mixed tussles and copulation attempts identified in CCHa2-R-RB^Gal4^ > Kir2.1 flies. (D) The temporal dynamic of anomaly scores during the mixed behavior, centralized at 1.67 s. SEM is indicated with the light color region. (E) Examples of close body contact behaviors identified in *Trh^Gal4^* flies. (F) The cosine similarity between the center frame of the close body contact behaviors (1.67s) and their local frames. SEM is indicated with the light color region. (G) The kicking index of *Trh^Gal4^* flies (n=30) was significantly lower than *w^1118^* flies (n=27), p = 0.0034, Mann Whitney test. (H) Examples of social aggregation behaviors of *Trh^Gal4^* flies and *w^1118^* flies. (I) Social distances of *Trh^Gal4^* flies (n=6) and *w^1118^* flies (n=6). *Trh^Gal4^* flies had much closer social distances with each other compared with *w^1118^* flies; nearest, p = 0.0043; median, p = 0.002; average, p = 0.0087; all Mann Whitney test. (J) Distributions of the median social distance of *Trh^Gal4^* flies and *w^1118^* flies. Distributions were calculated within each replication. Average distributions are indicated with solid lines, and SEMs are indicated with light color regions.

Some false positives occurred when only the IES was used as the anomaly score (Figure 3—figure supplement 1A). The reason could be that two flies in a chamber could be in mathematically infinite relative positions and form a vast event space. However, each group usually only contains several videos, and each video is only recorded for several minutes. For some rare postures, even though the probability of observing them is similar in both the query and reference group, they might only occur in the query group but not in the reference group. Therefore, an intra-group score (IAS) is introduced in the second step to eliminate these false-positive effects. We assume that those rare events should not be sampled frequently in the query groups either. Thus, the IAS is defined as the k-nearest distance of the query frame against all other frames within its group, except those within the time window of ±50 frames (Figure 3—figure supplement 1B). The final anomaly score is defined as the IES minus the IAS.

To test whether our methods could detect anomalous behavior in real-world data, we performed genetic screenings within fifteen neurotransmitter-related mutant alleles or neuron-silenced lines (with UAS-Kir2.1^39^) (Figure 3B). Their male-male interaction videos were inferred by Selfee trained on male-female courtship videos. Since we aimed to find interactions distinct from male-male courtship behaviors, a baseline of ppk23>Kir2.1 flies was established because this line exhibit strong male-male courtship behaviors^40^. We compared the top-100 anomaly scores from sets of videos from experimental groups and wild-type control flies. The results revealed that one line, CCHa2-R-RB>Kir2.1, showed a significantly high anomaly score. By manually going through all anomalous live-frames, we further identified its phenotype as a brief tussle behavior mixed with copulation attempts (Figure 3C, Video 1, 0.2x play speed). This behavior was ultra-fast and lasts for less than a quarter second (Figure 3D), making it difficult to be detected by human observers. Up to this point, we have demonstrated that the frame-level anomaly detection could capture sub-second behavior episode that human observers tend to neglect.

Selfee also revealed that *Trh* knock-out flies had an unusual close body contact during the screening. *Trh* is the crucial enzyme for serotonin biosynthesis, and its mutant flies showed a statistically significantly higher anomaly score (Figure 3B) than the wild-type control. Selfee identified 60 frames of abnormal behaviors within 42,000 input frames, occupying less than 0.15% of the total recording time. By manually going through all these frames, we concluded most of them as short-range body interactions (Figure 3E and Video 2, 0.2x play speed), and these social interactions could last for around half to one second on average (Figure 3F). Despite that serotonin signals were well-studied for controlling aggression behavior in flies^41^, to the best of our knowledge, the close body contact of flies and serotonergic neurons’ role in this behavior has not been reported yet. A possible reason is that this behavior has no unique posture compared with other behaviors, like wing extension, and this behavior is too scarce to be noticed by human experts.

To further ask whether these close body contacts have biological significance, we performed corresponded behavior assays on mutant flies. Based on the fact that the *Trh* mutant male flies have a higher tolerance to body touch, we hypothesized that they would have a decreased defensive behavior. As previously reported, fruit flies show robust defensive behavior to mechanical stimuli on their wings^42,43^. Decapitated flies would kick with their hind legs when a thin probe stimulates their wings. This stimulation mimics the invasion of parasitic mites and could be used to test its defensive behavior. Our results showed that *Trh* knock-out flies had a significantly lower kicking rate than control flies (Figure 3G), indicating a reduction of self-defensive intensity. Next, we performed social behavior assay^44,45^ on the mutant flies because the close body contact can also be explained by reduced social repulsion. We measured the nearest distance, median distance, and average distance of each male flies in a forty- individual group placed in a vertical triangular chamber. By comparing median values of these distances of each replication, *Trh* knock-out flies kept significantly shorter distances from others than the control group (Figure 3H, I). The probability density function of their median distances also showed that knock-out flies had a closer social distance than control flies (Figure 3J). Therefore, we concluded that *Trh* knock-out flies had reduced social repulsion. Taken together, Selfee is capable of discovering novel features of animal behaviors with biological relevance when a proper baseline is defined.

### Modeling motion structure of *Drosophila* courtship behaviors

Animal behaviors have long-term structures beyond single-frame postures. The duration and proportions of each bout and transition probabilities of different behaviors have been proven to have biological significance^18,46^. To better understand those long-term characteristics, we introduce AR-HMM and DTW analyses to model the temporal structure of *Drosophila* courtship behavior. AR-HMM is a powerful method to analyze stereotyped behavioral data^18,47,48^. It discovers modules of behaviors and describes the modules with auto-regressive matrixes. The transition probability of each state is defined by the transition matrix of the HMM (Figure 4A). In this way, AR-HMM could capture local structures of animal behaviors as well as syntaxes.

**Figure 4.**
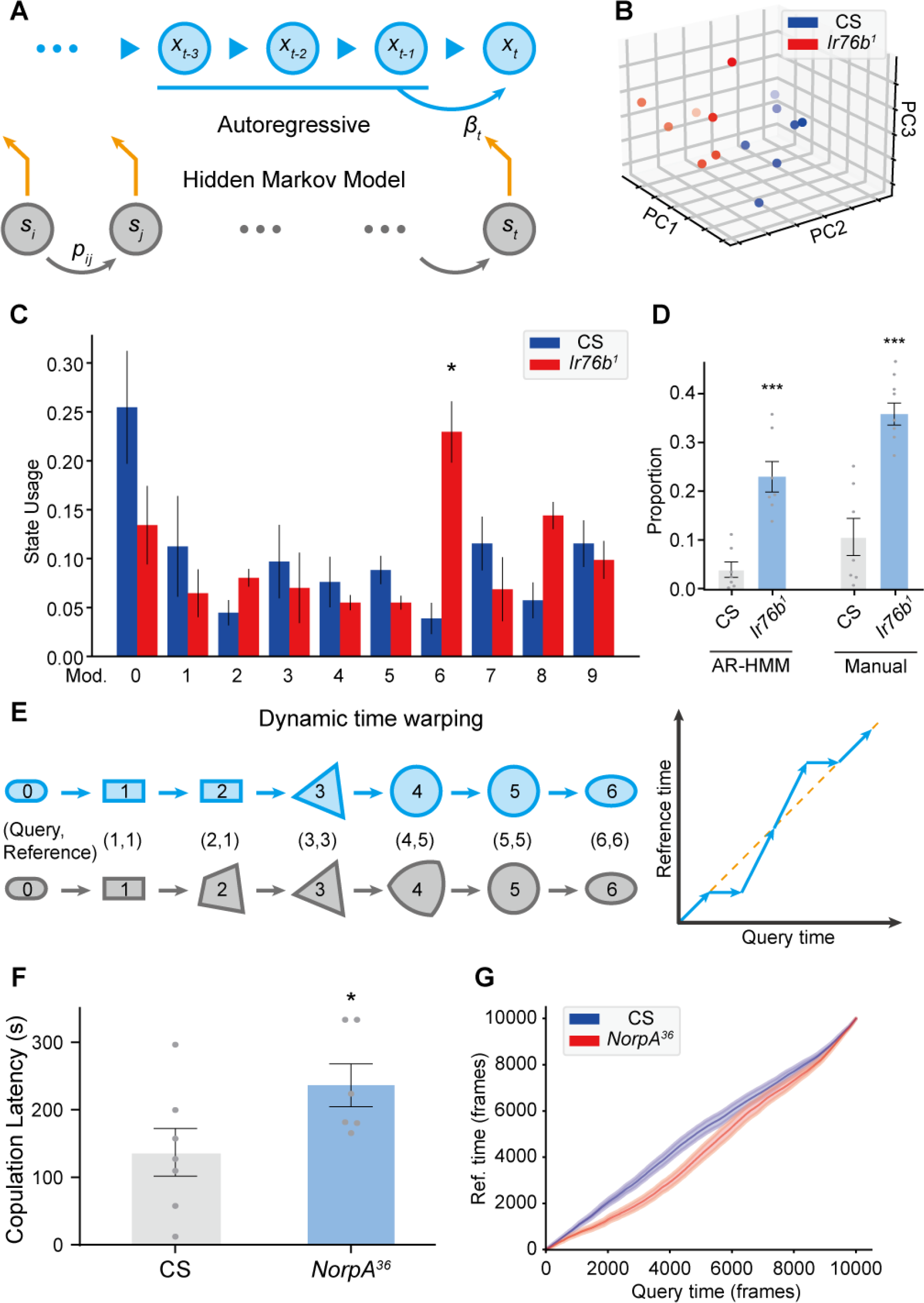
Time-series analyses using Selfee produced features. (A) A brief illustration of the AR-HMM model. The local autoregressive property is determined by *βt*, the autoregressive matrix, which is yield based on the current hidden state of the HMM. The transition between each hidden state is described by the transition matrix (*pij*). (B) PCA visualization of state usages of wild-type flies (n=7) and *Ir76b^1^* mutant flies (n=7). (C) State usages of ten modules. Module No. 6 showed significantly different usages in wild-type and mutant flies; q = 0.022, Mann Whitney test with Benjamini and Hochberg correction. (D) Module No.6 showed similar statistic results as manually labeled non-interactive behaviors. Module No.6, p = 0.0006; manually labeled non-interactive behaviors, p = 0.0006; Both were Mann Whitney test. (E) A brief illustration of the DTW model. The transformation from a rounded rectangle to an ellipse could contain six steps (grey reference shapes). The query transformation lags at step 2 but surpasses at step 4. The dynamic is visualized on the right panel. (F) *NorpA^36^* flies (n=6) showed a significantly longer copulation latency than wild-type flies (n=7), p = 0.0495, Mann Whitney test. (G) *NorpA^36^* flies had delayed courtship dynamics than wild-type flies with DTW visualization.

We asked if we could detect the dynamic changes of courtship behaviors of male flies by disturbing their chemosensation. *Ir76b* is an extensively studied (co)receptor that is known to mediate female pheromones detection^49-52^. We used an AR-HMM model with ten modules (No.1 to 10) to analyze the courtship of *Ir76b* mutant flies and their control group and focused on state usages. PCA of state usages revealed an apparent difference between mutant flies and control flies (Figure 4B). Module No.6 showed a statistically significant difference among ten discovered modules (Figure 4C). By manually going through all the frames of module No.6, we found that it mainly contained non-interactive behaviors with minor contaminations of courtship behaviors (Video 3, 1x play speed). To validate this result, we compared it with human annotations. Although this module did not cover all non-interactive behaviors that human experts would label, they showed a similar trend between the experimental and control group (Figure 4D). We also performed AR-HMM analysis with a much larger module number. The PCA result was also distinct, and the previous module No.6 was split into five smaller modules (No.2, 15, 24, 32, 34) containing non-interactive behaviors (Figure 4—figure supplement 1, Video 4-8, 1x play speed). This tuning indicated that AR-HMM analysis is robust regardless of the number of modules, same as a previous report^18^. Our results indicated that *Ir76b* mutation might affect male files’ detection of female pheromones and consequentially the temporal structure of their courtship behaviors. These findings prove that Selfee with AR-HMM could discover the differences in proportions of behaviors, similar to what was achieved with classic manual analysis such as the courtship index.

The AR-HMM modeling does not necessarily capture the difference of long-term dynamics intuitively, such as the latency of certain behaviors. To solve this problem, we introduce DTW analysis. DTW is a well-known algorithm to align time series, which returns the best-matched path and the matching similarity (Figure 4E). The alignment can be simplified as follows. When given the same start state and end state, it optimally maps all indices from the query series to the reference series monotonically. Pairs of mapped indices form a path to visualize the dynamic difference. The points upper than the diagonal line indicate that the current time point in the query group is matched to a future time point in the reference group so that the query group has faster dynamics and vice versa. In our experiments, cosine similarities of Selfee extracted representations are used to calculate warping paths.

Previously, DTW was widely applied to numerical measures of animal behaviors, including trajectory^53^, audios^54^, and acceleration^55^. For the first time, we applied DTW to image data, with the aid of Selfee, to study the prolonged dynamic of animal behaviors. We analyzed whether the vision is essential for a male fly’s copulation completion. Visual cues are essential for male flies to locate female flies during courtship^56^, and mutant flies of *NorpA,* which have defective visual transduction^57^, have a prolonged courtship latency in our experiments (Figure 4F), similar to previously findings^58^. When wild-type flies were used as the reference for the DTW, the group of *NorpA* mutant flies yielded a curve lower than the diagonal line, indicating a delay of their courtship behaviors (Figure 4G). In this way, our experiments confirm that Selfee and DTW could capture differences in long-term dynamics such as behavior latency. In conclusion, DTW and AR-HMM could capture temporal differences between control and experimental groups beyond single-frame postures, making Selfee a competent unsupervised method for traditional analyses like courtship index or copulation latency.

## DISCUSSION

Here we use cutting-edge self-supervised learning methods and convolutional neural networks to extract Meta-representations from animal behavior videos. Siamese CNNs have proven their capability to learn comprehensive representations^29^. The cosine similarity, part of its loss function used for training, is rational and well-suited to measure similarities between the raw images. Besides, convolutional neural networks are trained end-to-end so that preprocessing steps like segmentation or key points extraction is unnecessary. By incorporating Selfee with different post-processing methods, we can identify phenotypes of animal behaviors at different time scales. In the current work, we demonstrate that the extracted representations could be used not only for straightforward distance-based analyses such as t-SNE visualization or k-NN anomaly detection but also for sophisticated post-processing methods like AR-HMM. These validations confirm that the extracted Meta-representations are meaningful and valuable.

By applying our method to mice mating behavior, we show that our Selfee out- performed some of the widely used pose-estimation methods in multi-animal behavior analysis. The famous DeepLabCut and similar methods could identify human-defined key points on animals. However, when animals of the same color are recorded at a compromised resolution and their body contacts are intensive, the current version of DeepLabCut could hardly extract useful features (Figure 1—figure supplement 1, Video 9). The reason is that it is extremely difficult to unambiguously label body parts like nose, ears and hips when two mice are close enough, a task challenging even for human experts. By contrast, Selfee could readily identify the frame as “intromission” (Figure 1—figure supplement 1) as human experts would do. These results show that our methods could capture global characteristics of behaviors like human experts, making it well-suited for processing multi-animal behavior videos, compared with pose-estimation methods.

We also demonstrate that the cutting-edge self-supervised learning model is accessible to biology labs. Our model can be trained on only one RTX 3090 GPU with a batch size of 256 within only 8 hours with the help of the newly proposed CLD loss function^38^ and other improvements (see Methods for further details). Furthermore, when the model pre-trained with mice videos was applied to rat behaviors, we were able to achieve a zero-shot classification of five major types of social behaviors (Figure 2—figure supplement 4). Although the F1 score was only 49.6%, it still captured the major differences between similar behaviors, such as allogrooming and social nose contact. Thus, we have demonstrated that self-supervised learning could be easily achieved with limited computation resources and a much shorter time and could be transferred to datasets that share similar visual characteristics.

Despite those advantages, there are some limitations of Selfee. First, because each live- frame only contains three raw frames, our model could not capture much information on the animal motion. It becomes more evident when Selfee is applied to highly dynamic behaviors such as mice mating behaviors. This can be overcome by increasing the computation because commonly used 3D convolution^59^ or spatial-temporal attention^60^ is good at dynamic information extraction but requires much more computational resources. Second, as previously reported, CNNs are highly vulnerable to image texture^61^. We observed that certain types of beddings of the behavior chamber could profoundly affect the performance of our neural networks (Figure 1—figure supplement 2), so in some cases, background removal is necessary (see Methods for further details). Lastly, Selfee could only use discriminative features within each batch, without any negative samples provided, so minor irrelevant differences could be amplified and cause inconsistent results (named mode-split). This mode split may increase variations of downstream analyses.

We can envision at least two possible future directions for Selfee. One is to optimize the backbone of neural networks to extract better features. Advanced self-supervised learning methods like DINO^33^ (with visual transformers, ViTs) could separate objects from the background and extract more explainable representations. Besides, by using ViTs, the neural network could be more robust against distractive textures^62^. At the same time, more temporal information can also be incorporated for a better understanding of motions. Combining these two, equipping ViTs with spatial-temporal attention could capture more temporal information.

Another direction will be explainable behavior forecasting for a deeper understanding of animal behaviors. For a long time, behavior forecasting has been a field with extensive investigations in which RNNs, LSTMs, or transformers are usually applied ^9,60,63^. However, most of these works use coordinates of key points as inputs. Therefore, the trained model might predominantly focus on spatial movement information and discover fewer behavioral syntaxes. By representation learning, spatial information is essentially condensed so that more syntaxes might be highlighted. Transformer models for forecasting could capture correlations between sub-series as well as long-term trends like seasonality^64^. These deep learning methods would provide behavioral neuroscientists powerful tools to identify behavior motifs and syntaxes that organize stereotyped motifs beyond the Markov property.

## ACKNOWLEDGEMENTS

We thank members of the Zhang lab for discussions. This work was supported by grants 31871059 and 32022029 from the National Natural Science Foundation of China, grant Z181100001518001 from the Beijing Municipal Science & Technology Commission, and a ‘Brain+X’ seed grant from the IDG/McGovern Institute for Brain Research at Tsinghua to W.Z.. W.Z. is supported by Chinese Institute for Brain Research, Beijing.W.Z. is an awardee of the Young Thousand Talent Program of China.

## AUTHOR CONTRIBUTIONS

Y.J. and J.H. coded the Selfee neural network and other accessory parts. Y.J., X.G. and S.L. performed animal experiments and analyzed data. W.Z. and X.X. supervised the project. Y.J. and W.Z. wrote the manuscript. All authors discussed and commented on the manuscript.

## DATA AND CODE AVAILABILITY STATEMENT

Major data used in this study were uploaded to Dryad. Data could be accessed via: https://datadryad.org/stash/share/BnIoOnaweOn2fc-sllSO0FhJJqduXQYaNu-KuPgz394 or its DOI 10.5061/dryad.brv15dvb8. We also shared our pretrained weights with Google Drive: https://drive.google.com/file/d/1A3U5guNEKA3Bi9H3QnfstZDEZ6aesqcR/view?usp=sharing. With the uploaded dataset and pretrained weights, our experiments could be replicated. However, due to its huge size and the limited internet service resources, we are currently not able to share our full training dataset. The full dataset is as large as 400GB, which is hard to upload to a public server and will be difficult for others users to download.

For training dataset, it would be available from the corresponding author upon reasonable request, and then we can discuss how to transfer such a big dataset. No project proposal is needed as long as the dataset is not used for any commercial purpose.

Our Python scripts could be accessed on GitHub: https://github.com/EBGU/Selfee

Other software used in our project include ImageJ(https://imagej.net/software/fiji/) and GraphPad Prism(https://www.graphpad.com/).

All data used to plot graphs and charts in the manuscript can be fully accessed on Dryad (DOI 10.5061/dryad.brv15dvb8).

## MATERIALS AND METHODS

### Fly stocks

All fly strains were maintained under a 12 h/12 h light/dark cycle at 25℃ and 60% humidity (PERCIVAL incubator). The following fly lines were acquired from Bloomington *Drosophila* Stock Center: *CCHa1^attP^* (84458), *CCHa1-R^attP^* (84459), *CCHa2^attP^* (84460), *CCHa2-R^attP^* (84461), CCHa2-R-RA^Gal4^ (84603), CCHa2-R-RB^Gal4^ (84604), *CNMa^attP^* (84485), *Oamb^attP^* (84555), *Dop2R^KO^* (84720), *DopEcR^Gal4^* (84717), *SerT^attP^* (84572), *Trh^Gal4^* (86146), *TK^attP^* (84579), *Ir76b^1^* (51309), *NorpA^36^* (9048), UAS-Kir2.1 (6596). *Tdc2^RO54^* was a gift from Dr. Yufeng Pan at Southeast University, China. Taotie-Gal4 was a gift from Dr. Yan Zhu at Institute of Biophysics, Chinese Academy of Sciences, China.

### Fly courtship behavior and male-male interaction

Virgin female flies were raised for 4∼6 days in fifteen-fly groups, and naïve male flies were kept in isolated vials for 8∼12 days. All behavioral experiments were done under 25 °C and 45%∼50% humidity. Flies were transferred into a customized chamber of 3 mm height and 10 mm diameter by a homemade aspirator. Fly behaviors were recorded using a stereoscopic microscope-mounted with a CCD camera (Basler ORBIS OY-A622f-DC) at the resolution of 1000×500 (for two chambers at the same time), or 640×480 (for individual chambers) and a frame rate of 30 Hz. Five types of behaviors were annotated manually, including “chasing” (a male fly follows a female fly), “wing extension” (a male fly extends unilateral wing and orientates to the female to sing courtship son, “copulation attempt” (a male fly bends its abdomen toward the genitalia of the female or the unstable state that male fly mounts on a female with its wings open), and “copulation” (male fly mounts on a female in a stable posture for several minutes).

### Fly defensive behavior assay

The kicking behavior was tested based on previously reported paradigms^1,2^. Briefly, flies were raised in groups for 3∼5 days. Flies were anesthetized on ice, and then male flies were decapitalized and transferred to 35 mm Petri dishes with damped filter paper on the bottom to keep the moisture. Flies were allowed to recover for around 30 minutes in the dishes. The probe for stimulation was homemade from a heat-melt yellow pipette tip, and the probe’s tip was 0.3 mm. Each side of flies’ wing margin was gently touched 5 times, and the kicking behavior was recorded manually. The statistical analysis was performed with the Mann Whitney test with GraphPad Prism Software.

### Social behavior assay for flies

The social distance was tested based on the previously reported method^3^. Briefly, flies were raised in groups for 3 days. Flies were anesthetized paralyzed on ice, and male flies were picked and transferred to new vials (around 40 flies per vial). Flies were allowed to recover for one day. The vertical triangular chambers were cleaned with 75% ethanol and dried with paper towels. After assembly, flies were transferred into the chambers by a homemade aspirator. The photos were taken after 20 min, and the positions of each fly were manually marked in ImageJ. The social distances were measured with the lateral sides of the chambers (16.72 cm) as references, and the median values of the nearest, median, and average distance of each replication are calculated. The statistical analysis was performed with the Mann Whitney test in GraphPad Prism Software.

### Mice mating behavior assay

Wild-type mice of C57BL/6J were purchased from Slac Laboratory Animal (Shanghai). Adult (8-24 weeks old) male mice were used for sexual behavior analysis. All animals were housed under a reversed 12 h/12 h light-dark cycle with water and food *ad libitum* in the animal facility at the Institute of Neuroscience, Shanghai, China. All experiments were approved by the Animal Care and Use Committee of the Institute of Neuroscience, Chinese Academy of Sciences, Shanghai, China (IACUC No. NA-016-2016).

Male mice were singly housed for at least 3 days prior to sexual behavioral tests. All tests were initiated at least 1 hr after lights were switched off. Behavioral assays were recorded using infrared cameras at the frame rate of 30 Hz. Female mice were surgically ovariectomized and supplemented with hormones to induce receptivity. Hormones were suspended in sterile sunflower Selfee oil (Sigma-Aldrich, S5007) and injected 10 mg (in 50 mL oil) and 5 mg (in 50 mL oil) of 17b-estradiol benzoate (Sigma-Aldrich, E8875) 48 h and 24 h preceding the test, respectively. On the day of the test, 50 mg of progesterone (Sigma-Aldrich, P0130; in 50 mL oil) was injected 4–6 h prior to the test. Male animals were adapted 10 min to behavioral testing rooms where a recording chamber equipped with video acquisition systems was located. A hormonal primed ovariectomized C57BL/6J female (OVX) was introduced to the home cage of male mice and videotaped for 30 min. Mating behavior tests were repeated three times with different OVX at least three days apart. Videos were manually scored using a custom-written MATLAB program. The following criteria were used for behavioral annotation: active nose contacts initiated by male mouse towards the female’s genitals, body area, faces were defined collectively as “social interest”; male mouse climbs the back of the female and moves the pelvis were defined as “mount”; Rhythmic pelvic movements after mount were defined as “intromission”; a body rigidity posture after final deep thrust were defined as “ejaculation”.

### Data preprocessing, augmentation and sampling

Fly behavior videos were decomposed into frames by FFmpeg, and only the first 10,000 frames of each video were preserved and resized into images with a resolution of 224×224. For model training of *Drosophila* courtship behavior, each video was manually checked to ensure successful copulations within 10,000 frames.

Mice behavior videos were decomposed into frames by FFmpeg, and only frames of the first 30 min of each video were preserved. Frames were then preprocessed with OpenCV^4^ in Python. Behavior chambers in each video were manually marked, segmented, and resized into images of a resolution of 256×192. For background removal, the average frame of each video was subtracted from each frame, and noises were removed by a threshold of 25 and the median filter with a kernel size of 5. Finally, the contrast was adjusted with histogram equalization.

For data augmentations, crop, rotation, flip, Turbo, and color jitter were applied. For a given frame, it formed a live-frame with its preceding and succeeding frames. For flies’ behavior video, three frames were successive, and for mice, the preceding or succeeding frame is one frame away from the current frame due to their slower dynamics^5^. Each live-frame was randomly cropped into a smaller version containing more than 49% (70%×70%) of the original image; then the image was randomly (clockwise or anticlockwise) rotated for an angle smaller than the acute angle formed by the diagonal line and the vertical line, then the image would be vertically flipped, horizontally flipped, and/or applied the Turbo lookup table^6^ at the probability of 50%, respectively; and finally, the brightness, contrast, saturation, and hue were randomly adjusted within 10% variation. Notably, since the Turbo transformation is designed for grayscale images, for a motion-colored RGB image, each channel was transformed individually. After Turbo transformation, their corresponded channels were composited to form a new image.

For fly data sampling, all images of all videos were randomly ranked, and each batch contained 256 images from different videos. For mice data sampling, all images of each video were randomly ranked, and each batch contained 256 images from the same video. This strategy was designed to eliminate the inconsistency of recording conditions of mice that was more severe than flies.

### Selfee neural network and its training

All training and inference were accomplished on a workstation with 128GB RAM, AMD Ryzen 7 5800x, and one NVIDIA GeForce RTX 3090. Selfee neural network was constructed based on publications and source codes of BYOL^7^, SimSiam^8^, and CLD^9^ with PyTorch^10^. In brief, the last layer of ResNet-50 was removed, and a 3-layer 2048- dimension MLP was added as the projector. Hidden layers of the projector were followed by batch normalization (BN) and ReLU activation, and the output layer only had BN. The predictor was constructed with a 2-layer bottleneck MLP with a 512- dimension hidden layer and a 2048-dimension output layer. The hidden layer but not the output layer of the predictor had BN and ReLU. As for the group discriminator for CLD loss, it had only one normalized fully connected layer that projected 2048- dimension output to 1024 dimensions, followed by a customized normalization layer that was described in the paper of CLD^9^.

The loss function of Selfee had two major parts. The first part was the negative cosine loss^7,8^, and the second part was the CLD loss^9^. For a batch of *n* samples, *Z*, *P*, *V* represented the output of projector, predictor, and group discriminator of the main branch, respectively; *Z’*, *P’*, *V’* represented the output of the reference branch; and *sg* as the stop-gradient operator. After k-means clustering of *V*, the centroids of k classes were given by *M*, and labels of each sample were provided in the one-hot form as *L*. The hyperparameter *θ* was 0.07, and *λ* was 2. The loss function was given by the following equations:

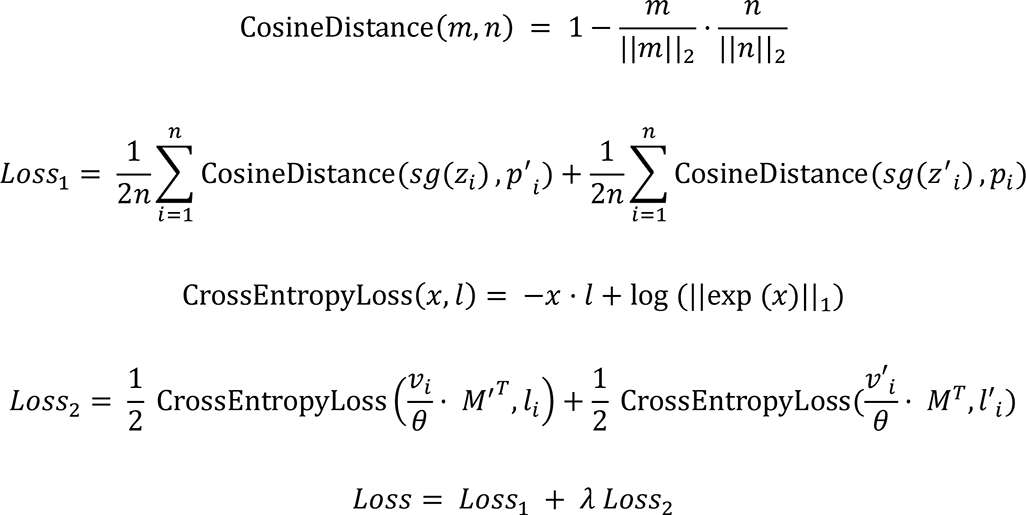

For all training processes, the Selfee network was trained for 20,000 steps with the SDG optimizer with a momentum of 0.9 and a weight decay of 1e-4. The learning rate was adjusted in the one-cycle learning rate policy^11^ with base learning rates and a pct start of 0.025. The model for *Drosophila* courtship behavior was initialized with ResNet-50 pre-trained on the ImageNet, and the base learning rate was 0.025 per batch size of 256. As for the mating behaviors of mice, the model was initialized with weights trained on the fly dataset, and the base learning rate was 0.05 per batch size of 256.

### t-SNE visualization

Video frames for t-SNE visualization were all processed by Selfee. Embeddings of three tandem frames were averaged to eliminate potential noises. All embeddings were transformed using t-SNE provided in the scikit-learn^12^ package in Python without further turning of parameters. Results were visualized with the Matplotlib^13^ package in Python, and their colors were assigned based on human annotations of video frames.

### Classification

Two kinds of classification methods were implemented, including the k-NN classifier and the LightGBM (Light Gradient Boosting Machine) classifier. The weighed k-NN classifier was constructed based on the previous reports^8,14^. LightGBM classifier^15^ was provided by its official package in Python. The F1 score and mAP were calculated with the scikit-learn^12^ package in Python.

For fly behavior classification, seven 10,000-frame videos were annotated manually. Seven-fold cross-validation was performed using embeddings generated by Selfee and the k-NN classifier. Inferred labels were forced to be continuous through time by using inferred labels of 21 neighbor frames to determine the final result. Then, a video independent of the cross-validation was annotated and inferred by a k-NN classifier using all 70,000 samples, and the last 3,000 frames were used for the raster plot.

For rat behavior classification, the RatSI dataset^16^ (a kind gift from Noldus Information Technology bv) contains nine manually annotated videos. We neglected three rarest annotated behaviors: moving away, nape attacking, and pinning, and we combined approaching and following into a larger category. Therefore, we used five kinds of behaviors, including allogrooming, approaching or following, social nose contact, solitary, and others. Nine-fold cross-validation was performed using embeddings generated by Selfee and the k-NN classifier. Inferred labels were forced to be continuous through time by using inferred labels of 81 neighbor frames to determine the final result.

For mice behavior classification, eight videos were annotated manually. Eight-fold cross-validation was performed using embeddings generated by Selfee and the k-NN classifier. To incorporate more temporal information, the LightGBM classifier and additional features were also used. Additional features include slide moving average and standard division of 81-frame time windows, the main frequencies, and their energy (using short-time Fourier transform in SciPy^17^) within 81-frame time windows. Early- stop was used to prevent over-fitting. Inferred labels were forced to be continuous through time by using inferred labels of 81 neighbor frames to determine the final result. Then, a video independent of the cross-validation was annotated and inferred by an ensemble classifier of eight previously constructed classifiers, and all frames were used for the raster plot.

### Anomaly detection

For a group of query embeddings of sequential frames *q1*, *q2*, *q3*, …, *qn*, and a group of reference embeddings of sequential frames *r1*, *r2*, *r3*, …, *rm*, the anomaly score of each query frame was given by the following equation:

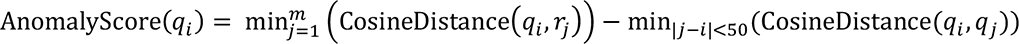

A PyTorch implementation of cosine similarity^18^ was used for accelerated calculations. The anomaly score of each video was the average anomaly score of the top 100 anomalous frames. The statistical analysis of the genetic screening was performed with one-way ANOVA with Benjamini and Hochberg correction in GraphPad Prism Software.

If negative controls are provided, anomalous frames are defined as frames with higher anomaly scores than the maximum anomaly score of frames in negative control videos.

### Autoregressive hidden Markov model (AR-HMM)

All AR-HMM models were built with the implementation of MoSeq^5^ (https://github.com/mattjj/pyhsmm-autoregressive). A principal component analysis (PCA) model that could explain 95% of variance of the control group was built and used to transform both control and experiment groups. The max module number was set as 10 for all experiments unless indicated otherwise. Each model was sampled for 1000 iterations. We kept other hyperparameters the same as the examples provided by this package. State usages of each module in control and experimental groups were analyzed by Mann Whitney test with SciPy^17^ followed with Benjamini and Hochberg correction. The state usages were also visualized after PCA dimensional reduction with scikit-learn^12^ and Matplotlib^13^.

### Dynamic time warping (DTW)

Dynamic time warping was modified from the Python implemention^19^ (https://dynamictimewarping.github.io/python/). Specifically, PyTorch implementation of cosine similarity^18^ was used for accelerated calculations.

### Pose-estimation with DeepLabCut

We used the official implementation of DeepLabCut^20,21^. For training, 120 frames of a mating behavior video were labeled manually, and 85% of them were used as the training set. Marked body parts included nose, ears, body center, hips, and bottom, following previous publications^22,23^. The model (ResNet-50 as the backbone) was trained for 100,000 iterations, with a batch size of 16. We kept other hyperparameters the same as default settings.

---

**Figure 1—figure supplement 1.**
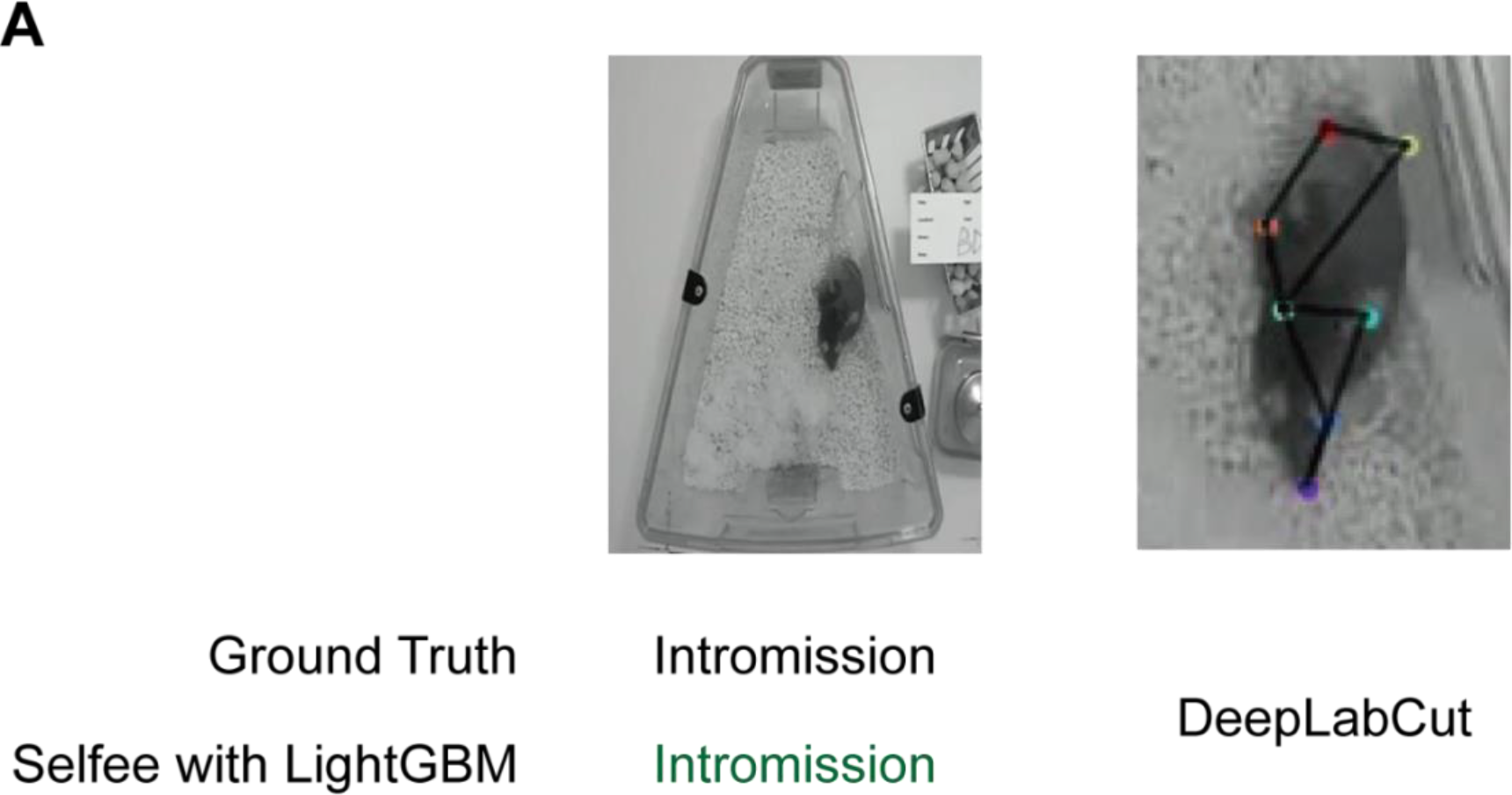
A comparison between Selfee and DeepLabCut on animals of the same color. (A) Selfee is more robust when applied to intensive social behaviors of animals of the same color, like mating behaviors of two black mice. The image is from the intromission behavior which could be identified by Selfee equipped with the trained LightGBM classifier. In contrast, a trained DeepLabCut model identified it as a single mouse.

**Figure 1—figure supplement 2.**
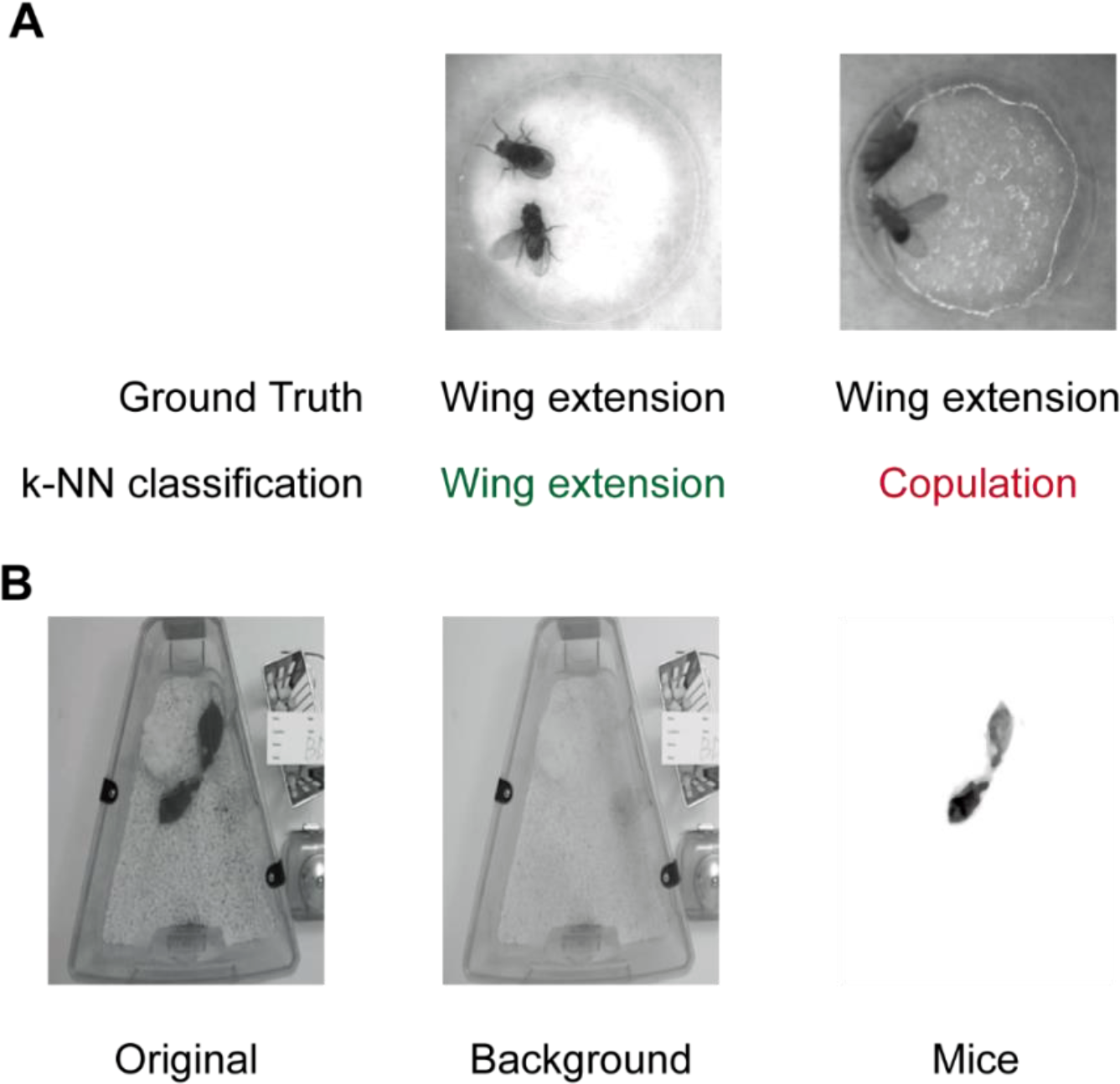
Beddings and backgrounds that affect training and inference of Selfee. (A) Textures on the damped filter paper would mislead Selfee to output features similar with copulation but not wing extension (ground truth). (B) Background inconsistency would affect the training process when Selfee was applied to mice behavior data. Therefore, backgrounds were removed from all frames to avoid potential defects.

**Figure 2—figure supplement 1.**
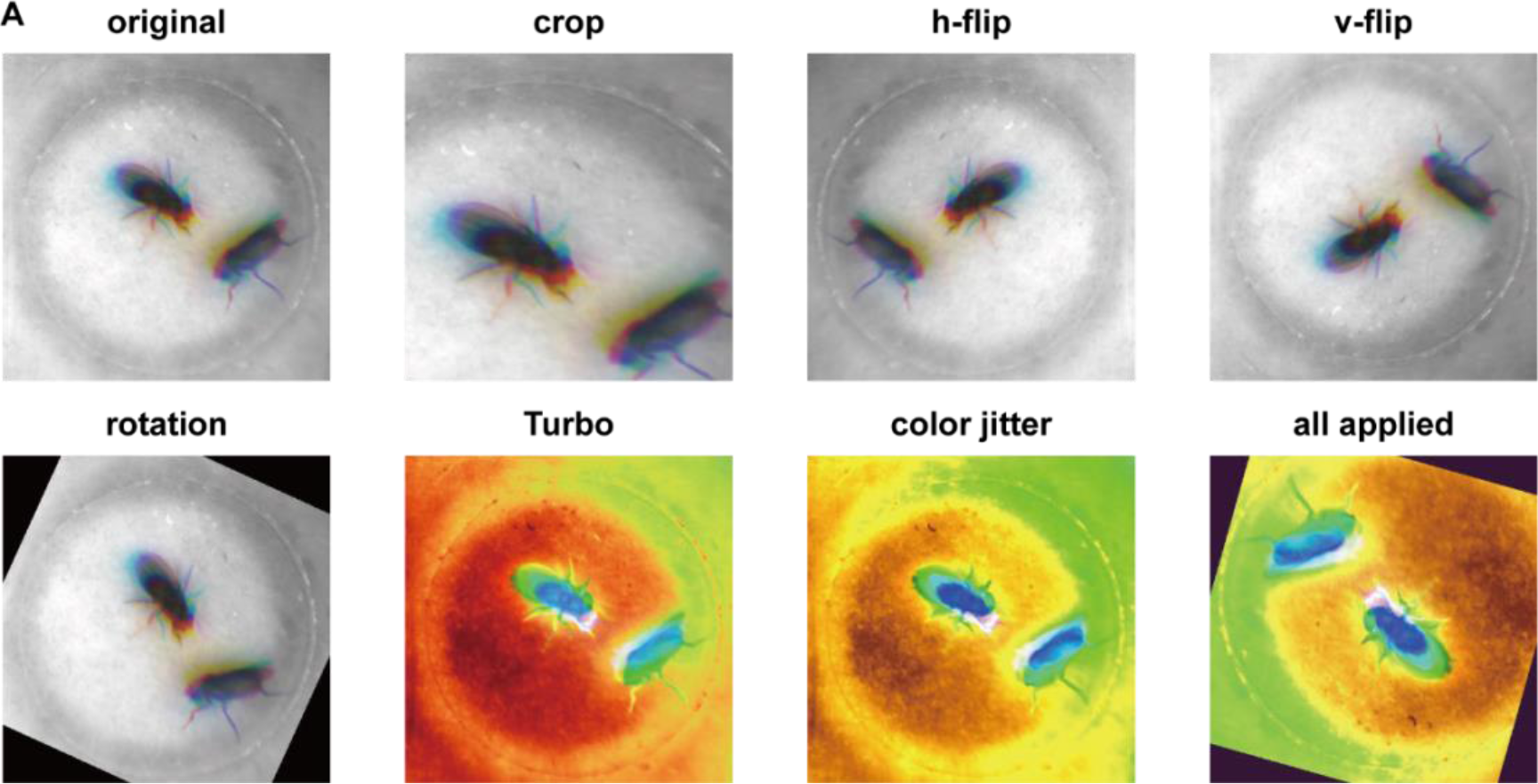
Different augmentations used for Selfee training. (A) Visualization of each augmentation. Detailed descriptions of each augmentation could be found in Methods and source codes.

**Figure 2—figure supplement 2.**
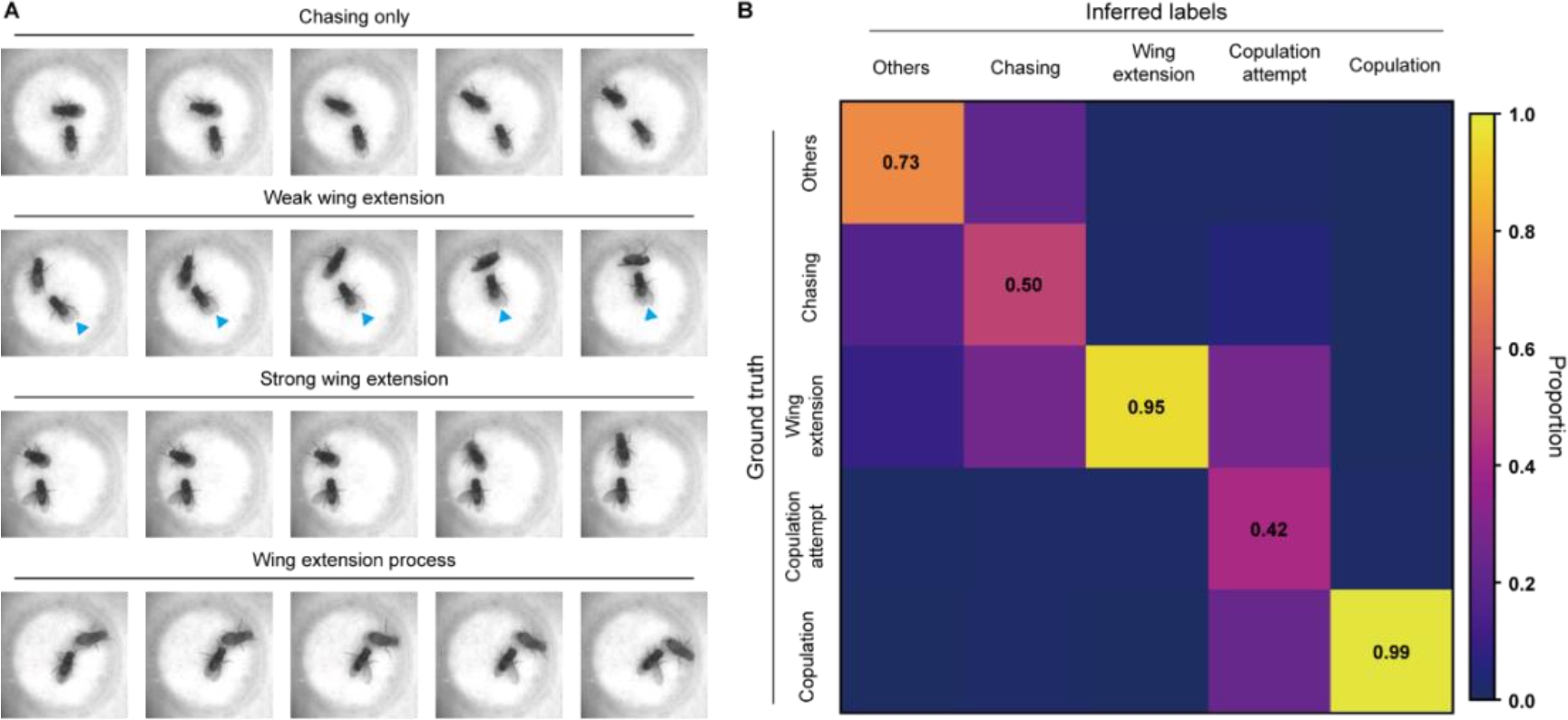
Difficulties on fly courtship behavior classification. (A) Some wing extension frames are hard to distinguish from chasing behaviors. Blue indicators point at slightly extended wings. (B) The confusion matrix of the k-NN classifier, normalized by the numbers of each behavior in inferred labels.

**Figure 2—figure supplement 3.**
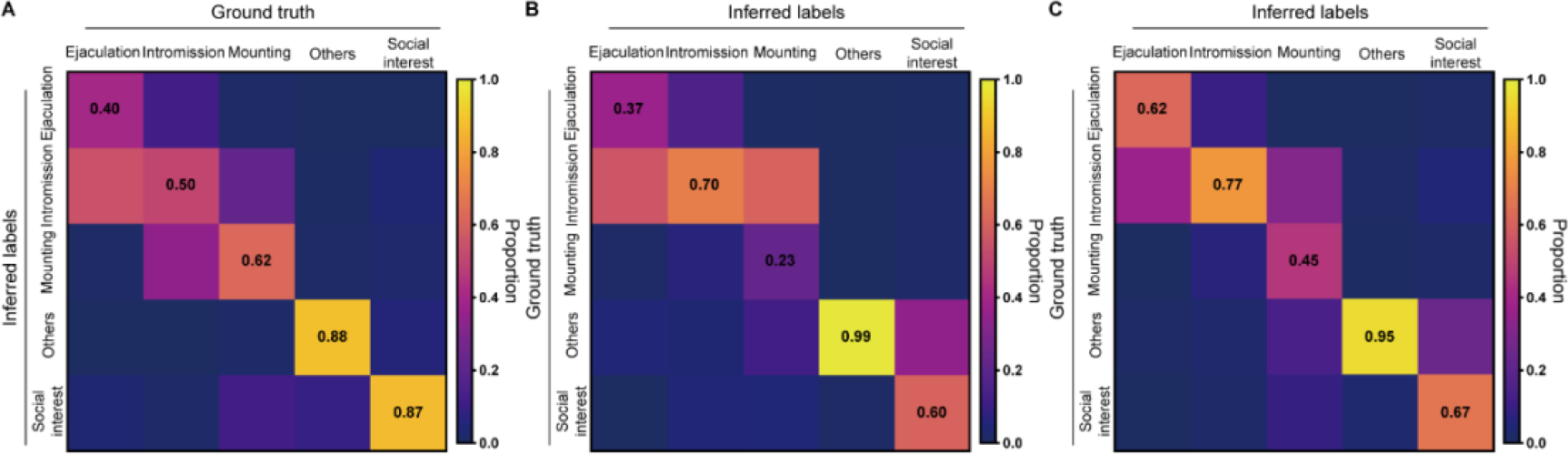
Classification of mice mating behaviors with Selfee extracted features. (A) For the k-NN classifier, the average F1 score of the nine-fold cross-validation was 59.0%, and mAP was 53.0%. The confusion matrix of the k-NN classifier, normalized by the numbers of each behavior in the ground truth. (B) The confusion matrix of the k-NN classifier, normalized by the numbers of each behavior in inferred labels. (C) The confusion matrix of the LightGBM classifier, normalized by the numbers of each behavior in inferred labels. The LightGBM classifier had a much better performance compared with the k-NN classifier.

**Figure 2—figure supplement 4.**
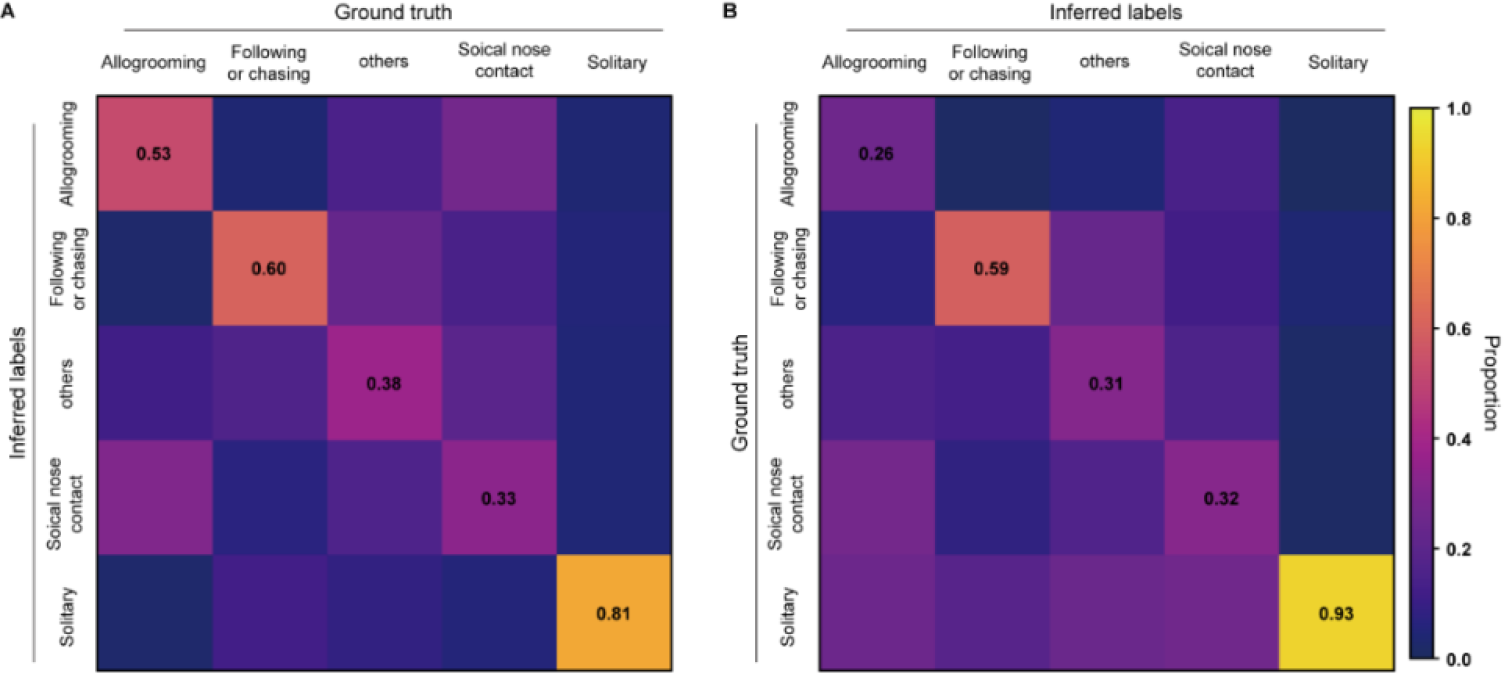
k-NN classification of rat behaviors with Selfee trained on mice datasets. (A) The average F1 score of the nine-fold cross-validation was 49.6%, and mAP was 46.6%. The confusion matrix of the k-NN classifier, normalized by the numbers of each behavior in the ground truth. (B) The confusion matrix of the k-NN classifier, normalized by the numbers of each behavior in inferred labels.

**Figure 2—figure supplement 5.**
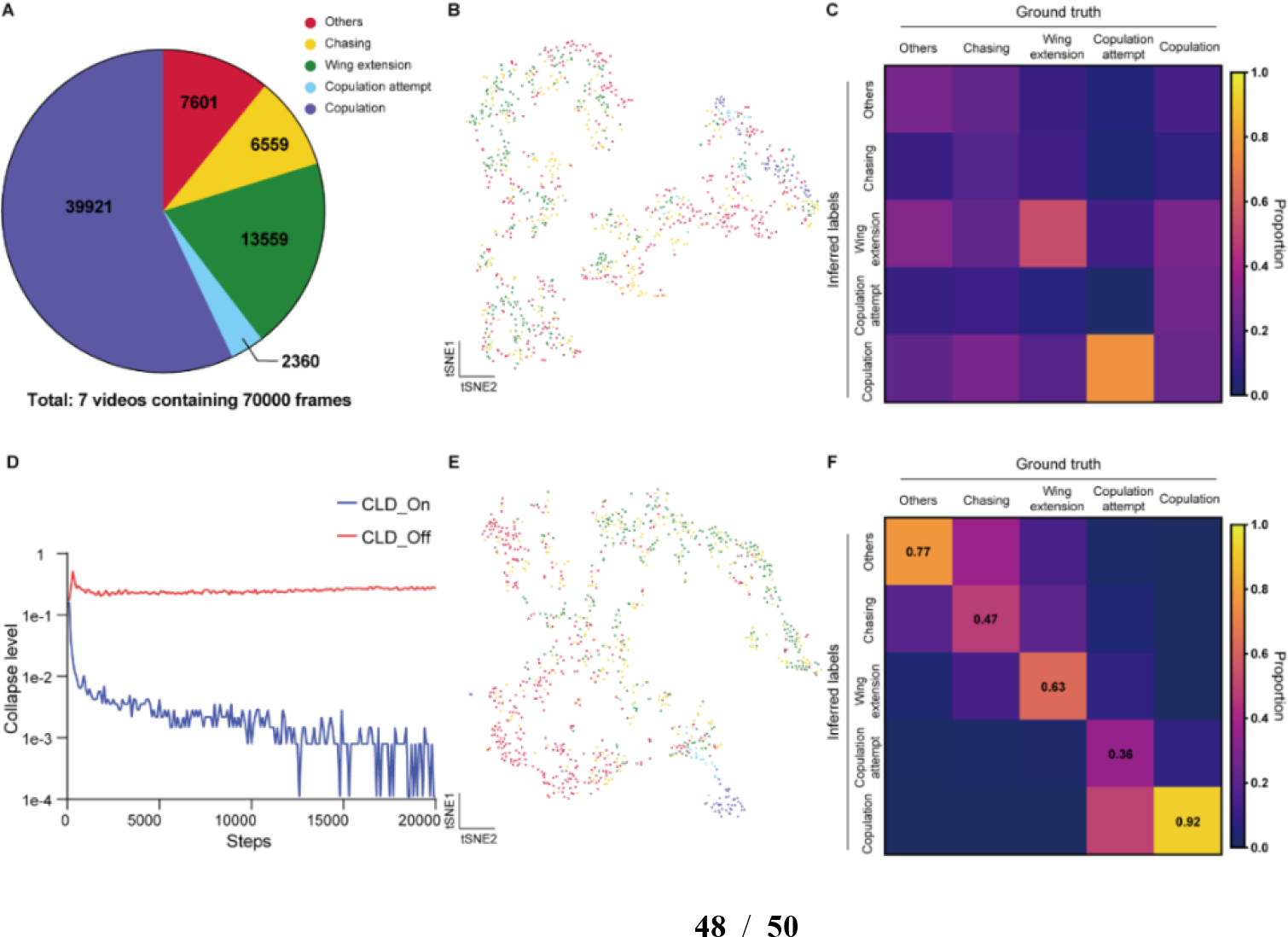
Ablation test of Selfee training process on fly datasets. (A) The distribution of different behaviors in wild-type flies courtship videos. (B) Visualization of the same live-frames as Figure 2D with t-SNE dimension reduction. Used representations were extracted by models trained without CLD loss. Each dot is colored based on human annotations. The legend is shared with panel A. (C) The confusion matrix of the k-NN classifier, normalized by the numbers of each behavior in the ground truth. Used representations were extracted by models trained without CLD loss. (D) Collapse levels during the training process. Without CLD loss, Selfee suffered from catastrophic mode collapse. Details for collapse level calculation could be found in Methods. (E) Visualization of the same live-frames as Figure 2D with t-SNE dimension reduction. Used representations were extracted by models trained without Turbo transformation. Each dot is colored based on human annotations. The legend is shared with panel A. (F) The confusion matrix of the k-NN classifier, normalized by the numbers of each behavior in the ground truth. Used representations were extracted by models trained without Turbo transformation.

**Figure 2—figure supplement 6.**
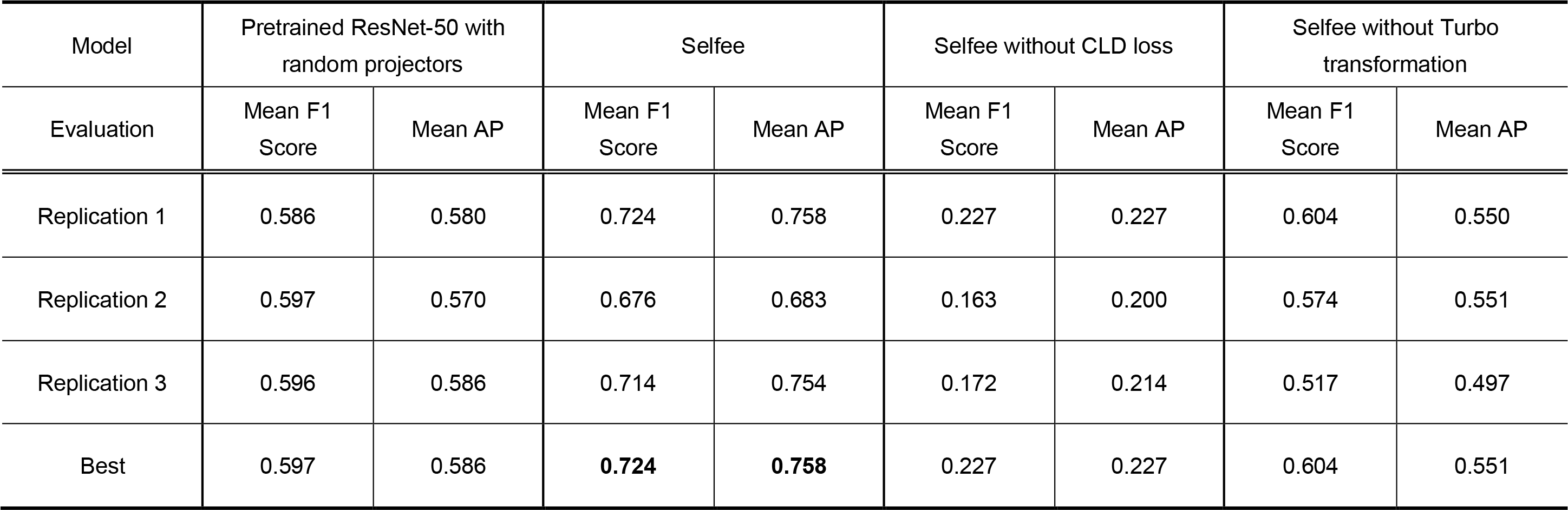
Ablation test of Selfee training process on fly datasets.

**Figure 3—figure supplement 1.**
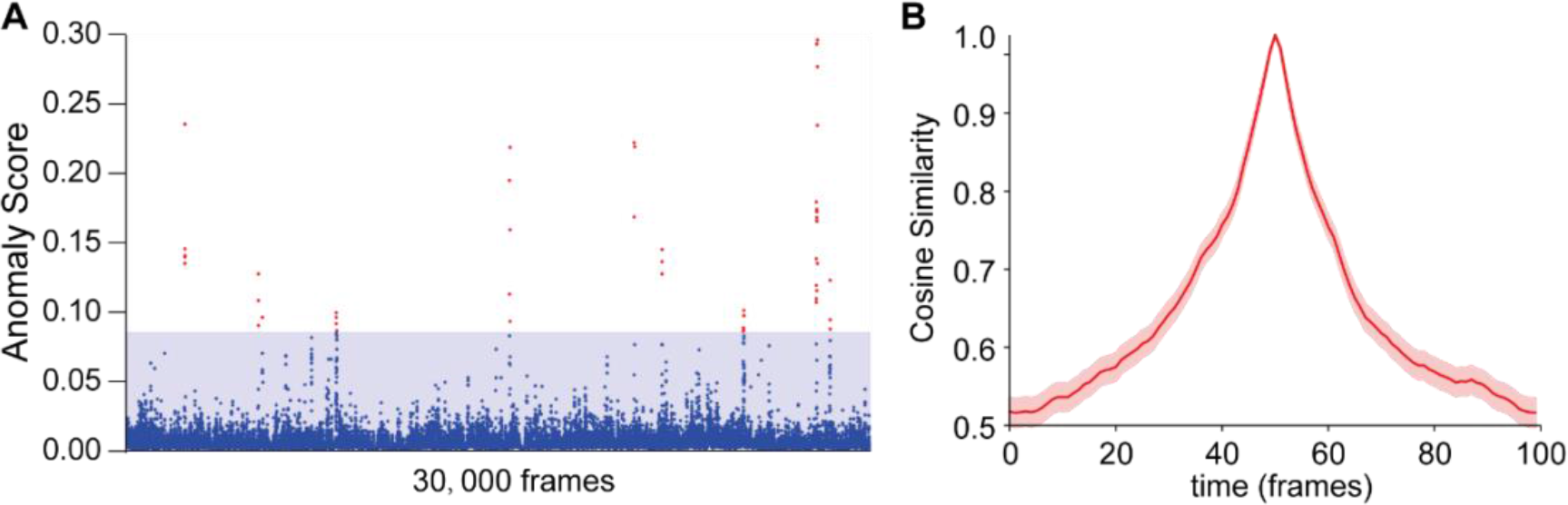
Using intra-group score (IAS) to eliminate false- positive results in anomaly detections. (A) Anomaly scores without IAS of wild-type male-male interactions with the same genotype as references. The blue region indicates the max anomaly score when using IAS; blue dots indicate anomaly scores without IAS that fall into the blue region; red dots indicate false-positive anomaly scores. (B) The cosine similarity between the center frame of wild-type courtship behaviors (1.67s) and their local frames. SEM is indicated with the light color region. Seven videos containing 70,000 frames were split into non-overlapping 100-frame fragments for calculations. Beyond ±50 frames, the cosine similarity droped to a much lower level, not affecting anomaly detection.

**Figure 4—figure supplement 1.**
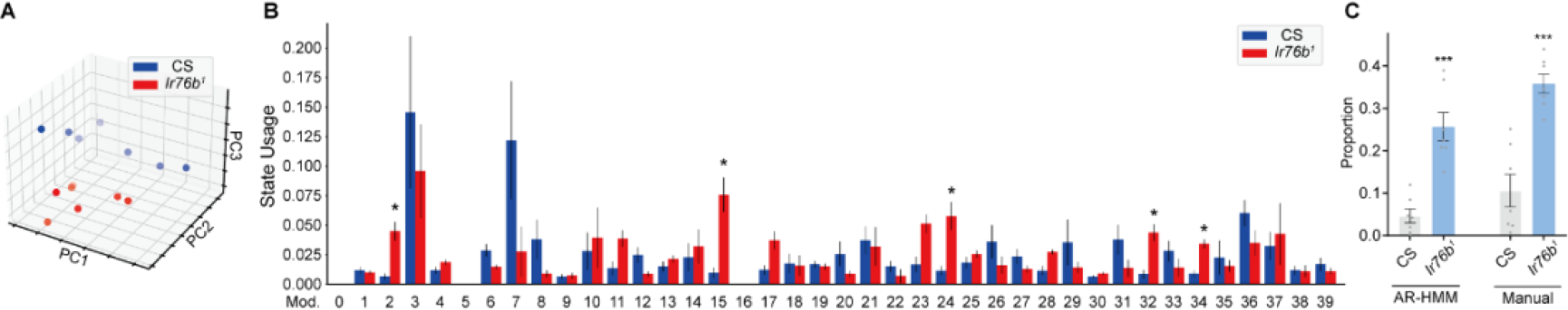
AR-HMM produced features with 40 modules using Selfee. (A) PCA visualization of state usages of wild-type flies and *Ir76b^1^* mutant flies. (B) State usages of forty modules. Module No. 2, 15, 24, 32, 34 showed significantly different usages in wild-type and mutant flies; q = 0.029, 0.029, 0.049, 0.049, 0.029 respectively, Mann Whitney test with Benjamini and Hochberg correction. (C) The collection of modules No. 2, 15, 24, 32, 34 showed similar statistic results as manually labeled non-interactive behaviors. AR-HMM module collection, p = 0.0006; manually labeled non-interactive behaviors, p = 0.0006; all Mann Whitney test.

